# Inter-strain variation in intra-chromosomal rates of recombination in *Caenorhabditis elegans*

**DOI:** 10.1101/2024.09.10.612278

**Authors:** Dharani U. Matharage, Anamica Khadgi, Amy L. Dapper

## Abstract

Meiotic recombination, the exchange of genetic material between homologous chromosomes, is a critical cellular process and a fundamental evolutionary parameter. Importantly, its rate varies dramatically across scales, from whole-genomes to kilobases. Although, the role of PRDM9 in intra-chromosomal variation in crossovers is well studied, our broader understanding of cellular and evolutionary dynamics driving intra-chromosomal patterns of recombination rate remains limited, in part due to the complexity of the landscape and the intrinsic difficulty of measuring this phenotype. Research on recombination rate variation in *Caenorhabditis elegans* is relatively sparse, but prior work suggests that this species provides a tractable system for studying intra-chromosomal recombination without many common confounding factors. Here, we measure variation in intra-chromosomal patterns of recombination rates in two genetically distinct populations (N2 and CB4856) of *C. elegans*. We find statistically significant, domain-specific differences in recombination rate between the two strains. Specifically, on chromosome IV, recombination is higher in the gene-rich central domain in N2, but higher in the gene-poor distal domain in CB4856. We detect no evidence of sex differences in recombination rate (heterochiasmy). Together, our findings demonstrate divergence in intra-chromosomal patterns of recombination between two widely utilized strains in a model system with an otherwise highly conserved recombination landscape.

## INTRODUCTION

Meiotic recombination occurs during the first stage of meiosis and results in the exchange of genetic material between homologous chromosomes. Variation in the rate of meiotic recombination has important cellular and evolutionary consequences. Deviations from an appropriate number and placement of crossovers can lead to errors in proper chromosomal segregation, resulting in inviable aneuploid gametes (Ferguson et al., 1996; Hassold and Hunt, 2001). By reducing linkage between loci, variation in recombination rate modulates the efficacy of selection and shapes major features of the genomic landscape, including nucleotide diversity and repetitive element density (Begun and Aquadro, 1992; Charlesworth et al., 1993; Comeron et al., 2012). As empirical approaches generate increasingly fine-scale maps of recombination rate variation across genomes (McVean et al., 2004; Kulathinal et al., 2008; Stevison et al., 2016; Beeson et al., 2019; i Torres et al., 2023), it is clear that in most species, chromosomal recombination landscapes are complex and dynamic. In some species, such as humans and mice, these landscapes are dominated by discrete, rapidly evolving recombination hotspots driven by PRDM9 (Ptak et al., 2005; Baudat et al., 2010; Paigen and Petkov, 2018). However, species with conserved hotspots, such as birds or yeast (Singhal et al., 2015; Lam and Keeney, 2015), and even species that appear to lack hotspots altogether, such as *Drosophila melanogaster* (Comeron et al., 2012), still exhibit dramatic fine-scale variation.

However, despite these advances, the evolutionary and cellular dynamics that shape these intra-chromosomal patterns remain poorly understood. Two major challenges limit advancements in this area: (1) the complexity of recombination landscape itself and (2) the intrinsic difficulty of measuring recombination rate. Most species exhibit fine-scale, broad-scale, and genome-wide variation in recombination rate, making it difficult to isolate one axis of variation (Coop et al., 2008; Chan et al., 2012; Dapper and Payseur, 2017). Furthermore, understanding intra-chromosomal variation from an evolutionary perspective requires measuring within population-level variation. This is difficult because it requires large-sample sizes that are difficult to obtain using widely-used methods, which are often time-consuming (i.e. pedigree analysis) and/or expensive (i.e. sperm sequencing), limiting the range of practical experimental approaches. Linkage disequilibrium-based methods circumvent some of these challenges, but provide population-averaged recombination rates, and are sensitive to assumptions about demographic history (Johnston and Cutler, 2012; Dapper and Payseur, 2018).

In contrast with other model organisms, *Caenorhabditis elegans* exhibits a uniquely simple recombination landscape. Genetic maps reveal that all five autosomes, as well as the X chromosome, are divided into five distinct domains of recombination (Rockman and Kruglyak, 2009): a central region with low rates of recombination, flanked by two distal regions with high rates of recombination and two small regions at each tip with zero recombination (Rockman and Kruglyak, 2009). Importantly, the transition in recombination rate between these regions is discrete and there is little evidence of fine-scale variation in recombination rate within each domain (Rockman and Kruglyak, 2009). Additionally, *C. elegans* show approximately complete crossover interference, such that each autosome and the X chromosome have exactly one crossover per meiosis (Hodgkin et al., 1979; Hillers and Villeneuve, 2003; Tsai et al., 2008), allowing intra-chromosomal variation in recombination rate to be disentangled from genome-wide variation. Gene density is also non-uniform across the chromosomes of *C. elegans*, providing an opportunity to test predictions about the relationship between gene-density and recombination rate. The large, central region with low recombination has high gene density (gene-rich center), while the two distal regions with high recombination have low gene density (gene-poor region) (Barnes et al., 1995).

Importantly, although *C. elegans* is predominantly selfing and exhibits low heterozygosity within strains (Barrie`re and Félix, 2005), this does not preclude selection on recombination rate. Meiotic recombination occurs during both spermatogenesis and oogenesis in hermaphrodites (Shakes et al., 2009; Cahoon et al., 2023). As such, genetic modifiers affecting crossover frequency or placement can influence fitness directly through their effects on chromosome segregation and indirectly through their effects on linkage among selected sites in both selfing and outcrossing individuals. This is supported by experimental evolution experiments in *C. elegans*, which provide evidence that recombination landscapes can evolve through indirect selection on a recombination modifier of large effect (*rec-1*), even in highly selfing populations (Parée and Teotónio, 2025). These results suggest that similar selective processes may shape naturally segregating differences in recombination rate observed between strains.

Because within-strain genetic diversity is limited, measuring intra-chromosomal recombination requires the introduction of heritable markers. Here, we utilize the inheritance patterns of fluorescent markers, inserted near the boundaries of the recombination domains on chromosome IV to estimate recombination rate within strains. This design enables direct comparison of recombination rates between two genetically distinct and widely used strains, N2 and CB4856, without relying on inter-strain crosses. The N2 strain is the reference wild-type strain and was isolated from mushroom compost in Bristol, England in 1951 (Brenner, 1974; Sterken et al., 2015). However, prior to its cryogenic preservation and dissemination, the N2 strain was maintained in the laboratory for many generations (from 1951 to 1969) allowing for the accumulation of adaptations to this environment (Sterken et al., 2015). As a result, the N2 strain exhibits many behavioral and physiological adaptations to a constant laboratory environment (De Bono and Bargmann, 1998; Styer et al., 2008; McGrath et al., 2009; Glauser et al., 2011; Andersen et al., 2014; Zhao et al., 2018) and higher fitness in the laboratory setting relative to wild isolates (Weber et al., 2010; Duveau and Félix, 2012). Importantly, unlike N2, CB4856, which was isolated from a pineapple field in 1972, did not undergo a long period of laboratory adaptation following its isolation (Hodgkin and Doniach, 1997; Volkers et al., 2013). These differences in evolutionary history provide a useful framework for examining divergence in recombination rate.

Previous work suggests that recombination rates in *C. elegans* may be flexible in response to variation in their physical and genomic environments, raising the prospect that there is standing variation for these traits in *C. elegans* under standard laboratory conditions. Recombination rates have been reported to vary with parental age and temperature in single-strain studies (Rose and Baillie, 1979), although other work finds little evidence for temperature effects (Rockman and Kruglyak, 2009). Evidence for sex differences in recombination rate (heterochiasmy) is also mixed, with some studies reporting higher recombination in hermaphrodites (Rose and Baillie, 1979), while others detect no significant differences (Lim et al., 2008).

Despite these insights, we do not have a clear picture of the degree to which recombination rate within these intra-chromosomal domains varies between strains of *C. elegans*. In this study, we estimate recombination rate in two adjacent chromosomal domains in two genetically-distinct strains. Importantly, we estimate recombination rate within each strain, rather than leveraging inter-strain crosses, allowing us to identify inter-strain variation. Specifically, we address the following questions: (1) Do genetically divergent strains of *C. elegans* differ in intra-chromosomal recombination rates in a constant laboratory environment? and (2) Do strains differ in the presence or magnitude of heterochiasmy?

## METHODS

### Strains

We measured recombination rate in two genetically distinct strains, N2 and CB4856, that represent well-characterized extremes in laboratory adaptation history and genome-wide genetic diversity (Sterken et al., 2015; Thompson et al., 2015). Both strains were obtained from the Caenorhabditis Genetics Center (CGC) (Stiernagle et al., 1999; Daul et al., 2019). We cultured the strains using standard methods and maintained them at 20C on NGM agar plates seeded with OP50 (Stiernagle, 2006). We performed all experiments at 20C.

### Introduction of the fluorescent markers

To estimate recombination rate, we utilized existing strains genetically engineered to carry fluorescent markers near the boundaries of the recombination domains on chromosome IV. The fluorescent marker strains (EG7910, EG8935, and EG7934) each carry a fluorescent marker (mCherry, GFP, and tdTomato respectively) in the N2 genetic background and were generated by Jorgensen lab and obtained from the CGC (Frøkjær-Jensen et al., 2014). These markers were inserted as single-copy transgenes using Mos1-mediated single-copy insertion (MosSCI), resulting in stable, defined genomic insertions that minimize position effects and copy-number variation (Frøkjær-Jensen et al., 2014). The marker positions on chromosome IV are 3,951,234 bp (mCherry), 13,023,274 bp (GFP), and 16,494,495 bp (tdTomato) (Frøkjær-Jensen et al., 2014).

The mCherry and GFP markers span the gene-rich center of the chromosome IV (approximately 3.8-12.9 Mb), which exhibits low recombination, while the GFP and tdTomato markers span a neighboring gene-poor distal domain with elevated recombination (approximately 12.9-16.7 Mb; (Rockman and Kruglyak, 2009). Markers were selected based on their position near the domain boundaries to maximize sensitivity to domain-specific recombination differences and due to their contrasting colors (red-green & green-orange) which makes their presence or absence easy to visualize in the same individual using the appropriate filters on a fluorescent microscope.

To generate strains with the desired genetic background (N2 or CB4856) and fluorescent markers on chromosome IV (GFP, tdTomato and mCherry), we backcrossed each marker strain to wild-type N2 and CB4856. The F1 offspring carrying the marker of interest were selected and backcrossed for nine additional generations, selecting for the marker each generation. After 10 rounds of backcrossing, we can expect 99.95% of the genome of each strain to be the desired genetic background and for each strain to be homozygous for the fluorescent marker (See Supplemental Methods). Because the fluorescent markers were introduced as defined single-copy insertions in the original marker strains, this backcrossing strategy also ensures that the identical insertion allele at the same genomic position is present in both genetic backgrounds, differing only in surrounding genetic context. Although linkage to the fluorescent insertions may retain small N2-derived genomic segments surrounding each marker in the CB4856 background, these segments are expected to be small and likely to reduce, rather than generate, between strain differences.

Utilizing this approach, we generated six new strains: three strains that each carry a single fluorescent marker (GFP, mCherry, or tdTomato) on a CB4856 genetic background and three strains that each carry a single fluorescent marker (GFP, mCherry, or tdTomato) on the N2 genetic background.

### Introduction of the fog-2 mutation

In wild-type strains, self-fertilization disrupts our ability to accurately estimate recombination rate by scoring inheritance of fluorescent markers. To eliminate selfing, we introduced the *fog-2* mutation into all six fluorescent strains. The *fog-2* mutation prevents hermaphrodites from producing viable self-fertilized embryos, ensuring that all offspring result from outcrossing (Schedl and Kimble, 1988).

We obtained strain JK574, which carries *fog-2* in the N2 background, from the CGC, and strain PTM299, which carries *fog-2* in the CB4856 background, from the McGrath lab at Georgia Institute of Technology. Using standard crossing schemes, we generated strains that were homozygous for the fluorescent marker of interest and the *fog-2* mutation in each genetic background (See Supplemental Methods). Because *fog-2* affects germline sex determination, not meiosis, and because it was present in all strains, it is unlikely to bias comparisons of recombination rate.

### Measuring Recombination Rate

To measure recombination rate, individuals from two homozygous fluorescent strains with the same genetic background (mCherry & GFP or GFP & tdTomato) were crossed to obtain offspring that are doubly heterozygous for both markers. We then crossed the resulting F1 progeny (focal individuals) with wild-type individuals, such that fluorescent markers could only be inherited from one parental individual. We performed this cross by placing five focal heterozygous hermaphrodites with five wild-type males of the same genetic background on an NGM plate. All adults were removed after 24 hours to synchronize offspring age. We scored the resulting offspring under a fluorescent scope once they reached the adult stage.

This cross produces four offspring phenotypes; two non-recombinant (single-marker phenotypes) and two recombinant (double-marker or no-marker phenotypes) (Supplemental Figure 1). Recombinant offspring frequencies were calculated as the proportion of total offspring displaying recombinant phenotypes. When scoring progeny, we found that GFP expression was sometimes difficult to identify reliably in males. To avoid errors in scoring, we excluded male offspring from recombination rate estimates and scored only hermaphrodite progeny. Because meiotic recombination occurs in the germline of the heterozygous parental individual, exclusion of male offspring should not bias recombination rate estimates. Multiple plates were scored per strain, sex, and chromosomal region, which was later incorporated into our statistical analyses. To assess heterochiasmy, we repeated the experimental design using doubly heterozygous males crossed to wild-type hermaphrodites, with all other procedures identical.

### Marker coverage of recombination domains

Because fluorescent markers were placed near—but not exactly at—recombination domain boundaries, some crossover events within domains may not have been captured. We estimated the proportion of recombination events missed based on domain sizes and observed recombination rates. These estimates assume approximately uniform recombination rates within domains, an expectation supported by the discrete and internally homogeneous recombination domains observed in *C. elegans* (Rockman and Kruglyak, 2009). These domain-wide estimates are presented for biological interpretation only; all statistical inference is based on the empirically observed marker intervals.

### Accounting for strain-specific differences in physical distance

Divergence in chromosome IV size between the N2 and CB4856 genomes, arising from insertions and deletions, could cause homologous genomic regions to differ in physical distance. Such differences could influence estimates of recombination frequency without reflecting evolutionary changes in the underlying recombination process. To evaluate whether observed strain-specific differences in recombination rate could arise from divergence in physical distance between marker pairs, we identified the precise insertion positions of each fluorescent marker in both N2 and CB4856 reference genomes. Flanking sequences were obtained from WormBuilder (Frøkjær-Jensen et al., 2014) and mapped using BLAST (Altschul et al., 1990). We calculated per-base-pair recombination rates and estimated the expected contribution of any divergence in physical distance to total recombination frequency.

### Statistical analysis

Recombination rates were analyzed using Bayesian generalized linear mixed models with a binomial error distribution and logit link, implemented in R using brms package (Bürkner, 2017, 2018, 2021). This framework is appropriate for recombination data, which consist of counts of recombinant versus non-recombinant offspring and follows a binomial sampling process. We included strain and sex as fixed effects to estimate biological differences in recombination probability, while plate was included as a random effect to account for shared environment, brood-level heterogeneity, and non-independence among offspring derived from the same experimental cross. Because offspring cannot be assigned to individual parents within a cross, recombination probabilities were estimated at the level of pooled offspring within each plate. The mixed-effects structure of this model allows variation among plates, arising from differences in fertility, brood size, or other cross-specific factors, to be incorporated directly into the inference, avoiding pseudo-replication and underestimation of uncertainty that can arise from contingency-table approaches (i.e. chi-squared test).

## RESULTS

### Variation in Recombination Rate between N2 and CB4856 Strains in the Gene-Rich Center

We estimated recombination rate in the gene-rich central region of the chromosome IV (Figure 1) for the N2 and CB4856 strains using a Bayesian generalized linear mixed model (GLMM), with strains and sex included as the fixed effects and the experimental plate included as a random effect. The intercept (*β* = -2.55) corresponds to the reference group, N2 hermaphrodites, and estimates a baseline recombination rate of approximately 7.3% with a 95% credible interval that the recombination rate lies between 6.3% and 8.3% (log-odds 95% credible interval: -2.70 to -2.40; Table 1). Relative to this baseline, the model provides strong support for lower recombination in the CB4856 strain, with the posterior distribution of the strain effect entirely below zero (*β* = -0.29, 95% credible interval: -0.49 to -0.09; Table 1), corresponding to an estimated recombination rate of 5.5% in CB4856. This represents an approximately 25% reduction in the odds of observing a recombination event in CB4856 compared to N2 in this interval. In contrast, we did not observe evidence of for sex-specific differences in recombination rate within this region (*β* = -0.07, 95% credible interval: -0.28 to 0.14; Table 1), indicating similar recombination probabilities in hermaphrodites and males within both strains in the gene-rich central region of chromosome IV (Table 1).

**Figure 1.**
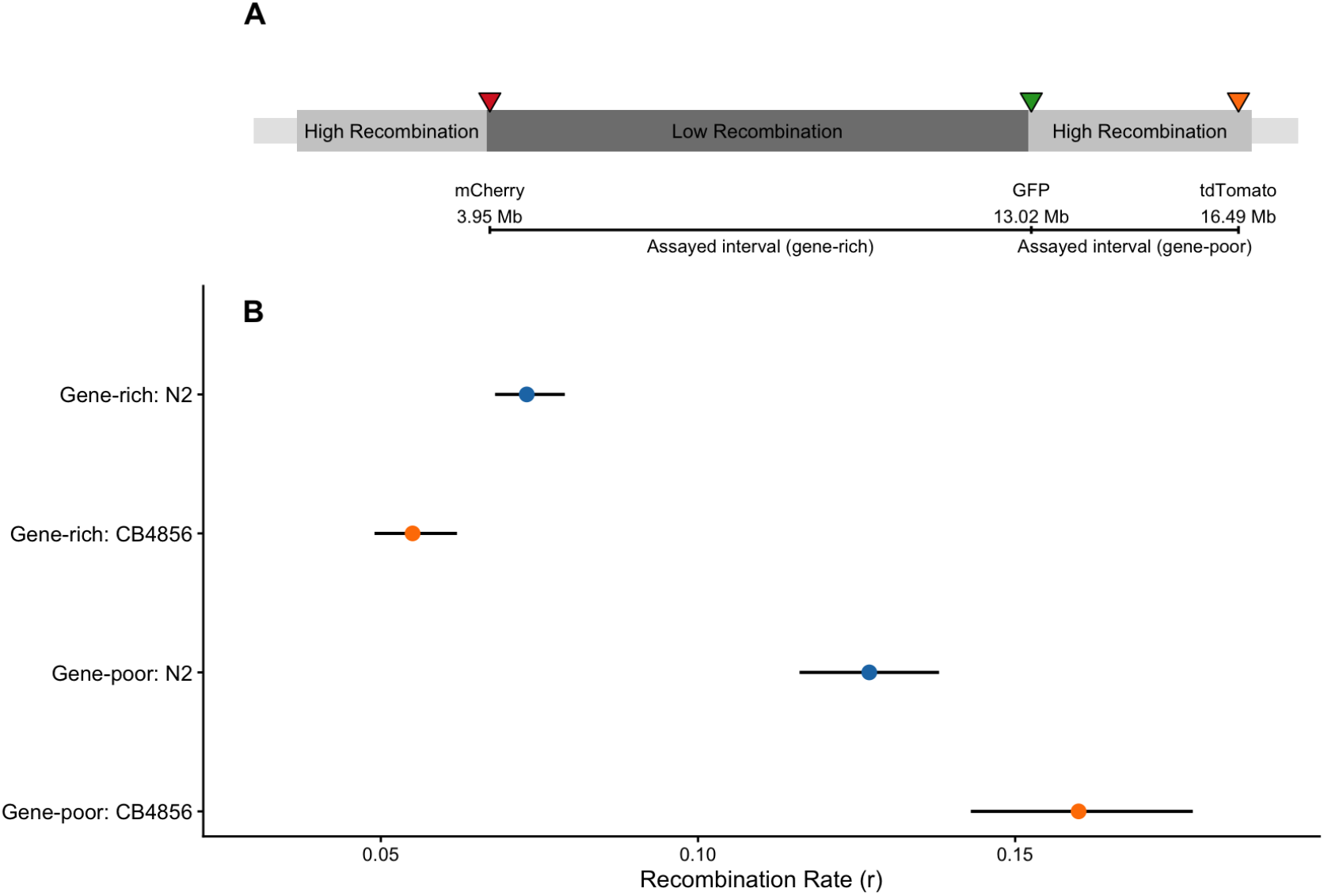
**(A)** Schematic of *C. elegans* chromosome IV showing large-scale recombination domains defined by genetic map estimates (Rockman and Kruglyak, 2009). Grey boxes indicate high-recombination, gene-poor distal arms flanking a low-recombination, gene-rich central domain (dark grey) and terminal regions with little to no recombination (light grey). Fluorescent markers (mCherry, GFP, tdTomato), inserted near domain boundaries, were used to assay recombination across a central gene-rich interval (mCherry–GFP) and a distal gene-poor interval (GFP–tdTomato). Marker positions are shown in megabases (Mb). **(B)** Posterior predicted recombinant offspring frequencies (*r*) from Bayesian generalized linear mixed models for each strain (N2, CB4856) across the two assayed intervals. Points show posterior mean estimates and horizontal bars indicate 1 posterior standard deviation (estimated error). Recombination differs between strains in a domain-specific manner; results are averaged across sexes, as sex-specific effects are minimal.

**Table 1.** Bayesian generalized linear mixed model estimates of recombinant offspring frequency (*r*) for gene-rich and gene-poor domains on chromosome IV. Estimates are on the log-odds scale. Credible intervals represent 95% posterior intervals.

| Domain | Term | Estimate | Est. Error | 2.5% | 97.5% |
| --- | --- | --- | --- | --- | --- |
| Gene-rich | Intercept | -2.548 | 0.079 | -2.705 | -2.396 |
|  | CB4856 | -0.291 | 0.104 | -0.492 | -0.086 |
|  | Male | -0.065 | 0.106 | -0.276 | 0.140 |
| Gene-poor | Intercept | -1.934 | 0.097 | -2.130 | -1.747 |
|  | CB4856 | 0.269 | 0.121 | 0.032 | 0.506 |
|  | Male | 0.060 | 0.129 | -0.202 | 0.316 |

### Variation in Recombination Rate between N2 and CB4856 Strains in the Gene-Poor Distal Region

We next assessed recombination in the adjacent gene-poor distal region of chromosome IV (Figure 1) using the same modeling framework. The intercept (*β* = -1.93) corresponds to the baseline recombination rate of the N2 hermaphrodites, and yields an estimated recombination probability of 12.7% with a 95% credible interval that recombination rate is between 10.6% and 14.8% (95% credible interval: -1.23 to -1.75; Table 1). Relative to this baseline, the model provides strong support that recombination rate is higher in the CB4856 strain in this interval, with the posterior distribution of the strain effect entirely above zero (*β* = 0.27, 95% credible interval: 0.03 to 0.51; Table 1). This effect corresponds to an increase in recombination probability from approximately 12.7% in N2 to 16% in CB4856, representing a 31% increase in the odds of a recombination event occurring in this interval. As in the gene-rich region, we found no evidence for sex-specific differences in recombination rate in the gene-poor region. The posterior distribution for the sex effect overlapped zero (*β* = 0.06, 95% credible interval: -0.20 to 0.32), indicating similar recombination probabilities in hermaphrodites and males in each strain (Table 1).

### Estimating recombination rates for the complete recombination domains

As the markers we used are located close to, but not exactly at the boundaries of recombination domain of interest on Chromosome IV, it is likely that we missed a small number of crossover events that occurred within the domain, but outside of the region covered by our markers. Given that recombination rates are expected to be roughly constant within domain, we can estimate the frequency of crossovers that were likely missed. The mCherry and GFP markers capture the chromosomal interval between 3.95Mb and 13.02Mb, while the gene-rich center is located between positions 3.90Mb and 12.97Mb (Rockman and Kruglyak, 2009), we estimate that we were unable to capture crossover events from the 0.05Mb of the domain at the mCherry marker’s end and captured additional crossover events for 0.05Mb that belong to the gene-poor distal region of the chromosome. Using the recombinant offspring frequency (Table 2), we estimated that we missed 0.04% of recombination events occurred in the neighboring domain and that we additionally captured 0.18% that occurred in the gene-poor region of the chromosome for the N2 strain. This gives us a total estimate of 6.96% for the entire gene-rich center in the N2 strain and, likewise, 5.23% for the CB4856 strain.

**Table 2.**
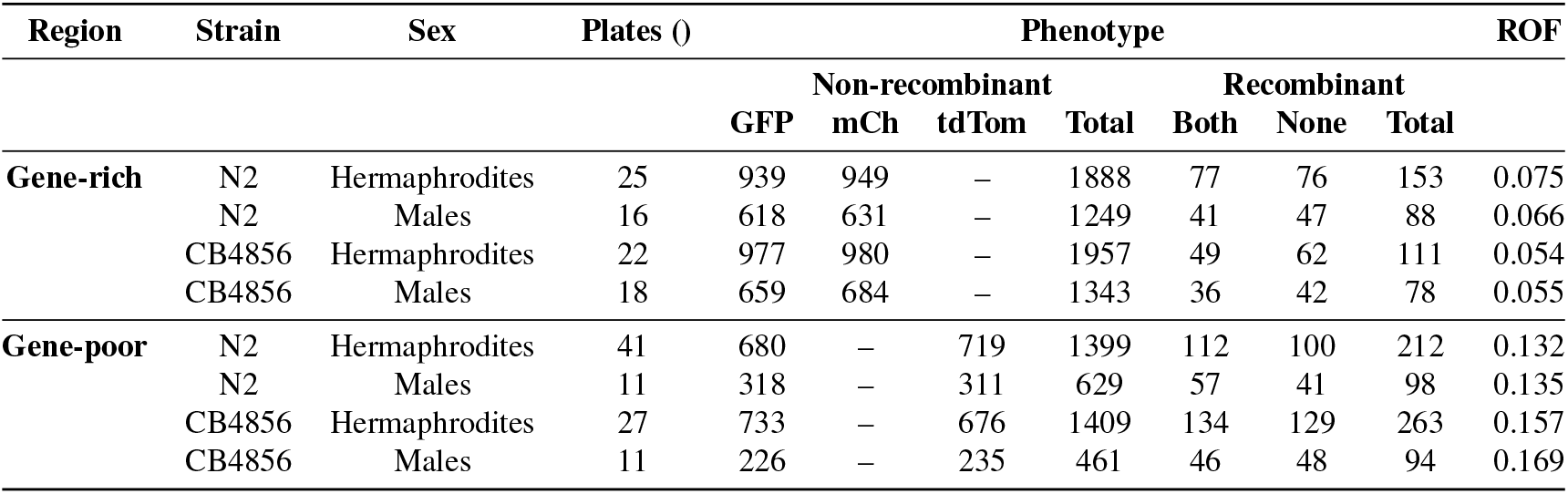
Counts of offspring phenotypes and recombinant offspring frequencies (ROF) for gene-rich and gene-poor regions of chromosome IV. Recombinant offspring frequency is calculated by dividing the number of recombinant offspring by the number of total offspring.

The GFP and tdTomato markers cover physical position 13.02Mb through 16.49Mb, while region with gene-poor region start at the physical position 12.97Mb and goes through 16.71Mb (Rockman and Kruglyak, 2009). Thus, we estimate that we were unable to capture recombination events over 0.27Mb of the domain. For the N2 strain, we observed a recombination rate of 13.3% across this 3.5Mb region (Table 2). Thus, we expect a recombination rate of 0.96% in the missing 0.27Mb, giving a total estimate of 14.26% for the entire gene-poor region in N2 strain. Likewise, for CB4856, we estimate that the missing segments have a recombination rate of 1.15%, bringing the total recombination rate of the domain to 17.15%.

We converted the recombination rate estimates in to cM/Mb, with the gene-rich region of the chromosome IV yielding a recombination density of 0.81 cM/Mb for the N2 strain and 0.61 cM/Mb for the CB4856 strain, where as the gene-poor region exhibited recombination densities of 3.4 cM/Mb for the N2 strain and 4.3 cM/Mb for the CB4856 strains. While not directly comparable due to the differences in cross setups, these estimates are generally consistent with prior estimates (Rockman and Kruglyak, 2009).

### Accounting for strain-specific differences in physical distance

Divergence in chromosome IV size between the N2 and CB4856 genomes, arising from insertions and deletions, could cause homologous genomic regions to differ in physical distance. Such differences could influence estimates of recombination frequency without reflecting evolutionary changes in the underlying cellular process. To evaluate whether the observed strain-specific differences in recombination rate could be explained by divergence in physical distance between marker pairs, we identified the precise genomic insertion positions of each fluorescent marker in both the N2 and CB4856 reference genomes. Using these positions, we calculated per-base-pair recombination rates and estimated the expected contribution of physical distance differences to total recombination frequency.

For the mCherry-GFP interval (gene-rich central region), the physical distance between the markers is 170,855 bp longer in CB4856 (N2: 9,072,063 bps, CB4856: 9,242,918 bps). Given a per base pair recombination rate in CB4856 of 6.0 × 10^*−*7^, this small expansion is expected to contributed 0.1% to the observed recombination rate. Correcting for this difference reduces the estimated effective recombination rate in CB4856 (5.4%) and therefore, increases the difference relative to N2 (7.1%) in this interval. Likewise, the physical distance between the tdTomato and GFP markers (gene-poor distal region) is 159,672 bp longer in CB4856 (N2: 3,471,201 bps, CB4856: 3,630,873). By calculating the per base pair recombination rate in CB4856 (4.3 × 10^*−*6^), we estimate that this expansion contributes 0.7% to the observed recombination rate in CB4856 (16.0%). While the additional physical distance accounts for a small portion of the strain effect in this interval, it cannot account for the majority of the difference compared to N2 (13.2%).

### No fitness effects from inserted fluorescent markers

We found no evidence that the fluorescent markers used to estimate recombination rate carried a fitness cost. Under Mendelian expectations, with no viability or scoring bias, the recombinant phenotypes (individual carrying both marker or neither) should occur in a 1:1 ratio. Consistent with this expectation, in both assayed intervals, recombinant offspring carrying both markers and those carrying neither marker were produced in approximately equal numbers (GFP - mCherry: 203 both vs. 227 none; p = 0. 2673, Exact binomial test; GFP - tdTomato: 349 both vs. 318 none; p = 0. 2454, Exact binomial test; Table 2). Likewise, non-recombinant single-marker phenotypes occurred at approximately equal frequencies in both intervals (GFP - mCherry: 3193 GFP vs. 3244 mCherry; p = 0.5332, Exact binomial test; GFP - tdTomato: 1957 GFP vs. 1941 tdTomato; p = 0.8101, Exact binomial test; Table2). The absence of significant deviations from expected 1:1 ratios for both recombinant and non-recombinant phenotypes indicates that early mortality or marker-associated fitness effects are unlikely to have influenced recombination rate estimates.

## DISCUSSION

By leveraging the discrete recombination domains that characterize the genomic landscape of *C. elegans*, we identified divergence in intra-chromosomal recombination rates between a laboratory-adapted strain (N2) and a wild-derived strain (CB4856). Specifically, we observed higher recombination in the gene-rich central domain of chromosome IV in the laboratory-adapted strain (N2), while the wild-derived strain (CB4856) exhibited higher recombination in the adjacent gene-poor distal domain. These results demonstrate that intra-chromosomal recombination rates can diverge between strains despite the highly conserved recombination landscape of *C. elegans*, which is characterized by complete crossover interference and minimal fine-scale variation (Rockman and Kruglyak, 2009).

Across both intervals examined on chromosome IV, we found no evidence of sex differences in recombination rate. The absence of heterochiasmy in these regions is consistent with prior work reporting similar recombination rates between hermaphrodites and males on chromosome III (Lim et al., 2008), but contrasts with studies that identified higher rates of recombination in hermaphrodites in other regions of the genome (Zetka and Rose, 1990; Meneely et al., 2002). Together these results suggest that heterochiasmy in *C. elegans* may be variable between chromosomes, regions, or strains.

While our study leverages a single pair-wise comparison of two recombination domains on one chromosome in two strains - variation in the evolutionary history of the strains, and the differences in gene density between the domains, provides an opportunity to consider potential evolutionary explanations for the observed patterns. One class of theoretical models predicts that increased recombination may be favored because it reduces linkage disequilibrium among loci under selection, thereby enhancing the efficacy of selection (Hill and Robertson, 1966; Felsenstein, 1974). Thus, higher recombination rates may be favored in gene-rich regions of the genome in populations experiencing variable or fluctuating environments because it accelerates adaptive evolution (Lenormand and Otto, 2000; Otto and Barton, 2001; Otto, 2009). However, the pattern we observe — higher recombination in the gene-rich region of the laboratory-adapted strain — does not align with this prediction.

An alternative interpretation is that reduced recombination is generally favored in gene-rich regions due to the fitness costs associated with disrupting co-adapted gene complexes (Felsenstein, 1974; Feldman et al., 1980; Barton, 1995). Under this scenario, divergence arises when selection maintaining low recombination is weakened. The prolonged maintenance of N2 under constant laboratory conditions (Andersen et al., 2012; Sterken et al., 2015), combined with repeated population bottlenecks and very low levels of genetic diversity (Barrie`re and Félix, 2005), may have relaxed selection on recombination modifiers, allowing recombination rates in gene-rich regions to increase through drift or weak selection. The strong and conserved suppression of recombination in the gene-dense centers of *C. elegans* chromosomes, a pattern that is conserved in related species, suggests that such costs may pervasive among this clade (Rockman and Kruglyak, 2009; Andersen et al., 2012; Noble et al., 2021; Teterina et al., 2023). However, we emphasize that this interpretation remains speculative and cannot be distinguished from alternative mechanisms with the current data.

Importantly, all recombination rates were estimated in individuals exposed to a constant laboratory environment. Yet, there is evidence that recombination rate may be plastic in *C. elegans*, varying in response to maternal age and temperature (Rose and Baillie, 1979; Lim et al., 2008). This raises the prospect that elevated recombination in gene-rich regions may be condition-dependent, with differences between the strains changing in direction or magnitude following exposure to biotic and/or abiotic stressors. The experimental framework developed here provides a foundation for testing this possibility in future studies.

Finally, it is important to consider the possibility that the observed divergence in recombination rate are driven by mechanisms not considered in our study, such as crossover assurance, or that it is not adaptive at all. Laboratory populations are, by necessity, finite and experience repeated bottlenecks during strain generation and maintenance, creating the opportunity for genetic drift to quickly generate phenotypic divergence. Yet, even if this divergence is neutral, it is significant, because it demonstrates that there is segregating variation in intra-chromosomal recombination rates within *C. elegans*, which have a notoriously conserved recombination landscape along other axes of variation. Here, our experimental design allows us to investigate between strain differences in the intra-chromosomal patterns of recombination, which not possible when utilizing inter-strain crosses. This result, coupled with the unique and simple recombination landscape and experimental tractability, supports the use of *C. elegans* as a powerful system to study genetic architecture and evolution of intra-chromosomal recombination landscapes.

## CONCLUSIONS

In conclusion, our findings demonstrate that intra-chromosomal recombination rates can differ between *C. elegans* strains despite the highly conserved nature of the *C. elegans* recombination landscape. Specifically, in chromosome IV, we clearly show that recombination rate was elevated within the gene-rich region in the laboratory-adapted strain whereas wild-derived strain exhibit higher recombination rates in the neighboring gene-poor region. We do not find any evidence of heterochiasmy in either focal region in either strain. Our result is significant because it raises the prospect that we can use *C. elegans* as a model system to dissect the genetic underpinnings of intra-chromosomal variation in recombination rate. An important next step will be to determine whether these strain-specific patterns are stable across environments, particularly following exposure to biotic or abiotic stress. The approach we have developed here provides a useful framework for addressing these questions and investigating the factors that shape recombination rate variation within species.

## ACKNOWLEDGMENTS

We thank Ahna Skop, Levi Morran, Janna Fierst, Chloe Girard, Bret Payseur, Jean-Francois Gout, Mark Welch, Donna Gordon and members of the Dapper lab and Gout lab for feedback on the experimental design and the interpretation of results. This work was supported by the U.S. National Science Foun-dation CAREER program (DEB-2143063) awarded to ALD. We also thank Brown lab for providing the fluorescent scope for nematode scoring. Some strains were provided by the CGC, which is funded by NIH Office of Research Infrastructure Programs (P40 OD010440). Portions of this manuscript were edited for language clarity using a large language model. All scientific content and interpretations are the responsibility of the authors.

## Notes

### Competing Interest Statement

The authors have declared no competing interest.

### Summary of Updates

This version includes minor revisions to improve the clarity, organization, and presentation of the manuscript, as well as updates to the statistical methods and analyses. No major changes were made to the study's findings or conclusions.

## REFERENCES

Altschul, S. F., Gish, W., Miller, W., Myers, E. W., and Lipman, D. J. (1990). Basic local alignment search tool. Journal of molecular biology, 215(3):403–410.

Andersen, E. C., Bloom, J. S., Gerke, J. P., and Kruglyak, L. (2014). A variant in the neuropeptide receptor npr-1 is a major determinant of caenorhabditis elegans growth and physiology. PLoS genetics, 10(2):e1004156.

Andersen, E. C., Gerke, J. P., Shapiro, J. A., Crissman, J. R., Ghosh, R., Bloom, J. S., Félix, M.-A., and Kruglyak, L. (2012). Chromosome-scale selective sweeps shape caenorhabditis elegans genomic diversity. Nature genetics, 44(3):285–290.

Barnes, T., Kohara, Y., Coulson, A., and Hekimi, S. (1995). Meiotic recombination, noncoding dna and genomic organization in caenorhabditis elegans. Genetics, 141(1):159–179.

Barrière, A. and Félix, M.-A. (2005). High local genetic diversity and low outcrossing rate in caenorhab-ditis elegans natural populations. Current Biology, 15(13):1176–1184.

Barton, N. H. (1995). A general model for the evolution of recombination. Genetics Research, 65(2):123– 144.

Baudat, F., Buard, J., Grey, C., Fledel-Alon, A., Ober, C., Przeworski, M., Coop, G., and De Massy, B. (2010). Prdm9 is a major determinant of meiotic recombination hotspots in humans and mice. Science, 327(5967):836–840.

Beeson, S. K., Mickelson, J. R., and McCue, M. E. (2019). Exploration of fine-scale recombination rate variation in the domestic horse. Genome research, 29(10):1744–1752.

Begun, D. J. and Aquadro, C. F. (1992). Levels of naturally occurring dna polymorphism correlate with recombination rates in d. melanogaster. Nature, 356(6369):519–520.

Brenner, S. (1974). The genetics of caenorhabditis elegans. Genetics, 77(1):71–94.

Bürkner, P.-C. (2017). brms: An r package for bayesian multilevel models using stan. Journal of statistical software, 80:1–28.

Bürkner, P.-C. (2018). Advanced bayesian multilevel modeling with the r package brms. the r journal, 10 (1), 395–411.

Bürkner, P.-C. (2021). Bayesian item response modeling in r with brms and stan. Journal of statistical software, 100:1–54.

Cahoon, C. K., Richter, C. M., Dayton, A. E., and Libuda, D. E. (2023). Sexual dimorphic regulation of recombination by the synaptonemal complex in c. elegans. Elife, 12:e84538.

Chan, A. H., Jenkins, P. A., and Song, Y. S. (2012). Genome-wide fine-scale recombination rate variation in drosophila melanogaster. PLoS genetics, 8(12):e1003090.

Charlesworth, B., Morgan, M., and Charlesworth, D. (1993). The effect of deleterious mutations on neutral molecular variation. Genetics, 134(4):1289–1303.

Comeron, J. M., Ratnappan, R., and Bailin, S. (2012). The many landscapes of recombination in drosophila melanogaster. PLoS Genetics, 8(10):e1002905.

Coop, G., Wen, X., Ober, C., Pritchard, J. K., and Przeworski, M. (2008). High-resolution mapping of crossovers reveals extensive variation in fine-scale recombination patterns among humans. science, 319(5868):1395–1398.

Dapper, A. L. and Payseur, B. A. (2017). Connecting theory and data to understand recombination rate evolution. Philosophical Transactions of the Royal Society B: Biological Sciences, 372(1736):20160469.

Dapper, A. L. and Payseur, B. A. (2018). Effects of demographic history on the detection of recombination hotspots from linkage disequilibrium. Molecular Biology and Evolution, 35(2):335–353.

Daul, A. L., Andersen, E. C., and Rougvie, A. E. (2019). The caenorhabditis genetics center (cgc) and the caenorhabditis elegans natural diversity resource. In The biological resources of model organisms, pages 69–94. CRC Press.

De Bono, M. and Bargmann, C. I. (1998). Natural variation in a neuropeptide y receptor homolog modifies social behavior and food response in c. elegans. Cell, 94(5):679–689.

Duveau, F. and Félix, M.-A. (2012). Role of pleiotropy in the evolution of a cryptic developmental variation in caenorhabditis elegans. PLoS biology, 10(1):e1001230.

Feldman, M. W., Christiansen, F. B., and Brooks, L. D. (1980). Evolution of recombination in a constant environment. Proceedings of the National Academy of Sciences, 77(8):4838–4841.

Felsenstein, J. (1974). The evolutionary advantage of recombination. Genetics, 78(2):737–756.

Ferguson, L. R., Allen, J. W., and Mason, J. M. (1996). Meiotic recombination and germ cell aneuploidy. Environmental and molecular mutagenesis, 28(3):192–210.

Frøkjær-Jensen, C., Davis, M. W., Sarov, M., Taylor, J., Flibotte, S., LaBella, M., Pozniakovsky, A., Mo-erman, D. G., and Jorgensen, E. M. (2014). Random and targeted transgene insertion in caenorhabditis elegans using a modified mos1 transposon. Nature methods, 11(5):529–534.

Glauser, D. A., Chen, W. C., Agin, R., MacInnis, B. L., Hellman, A. B., Garrity, P. A., Tan, M.-W., and Goodman, M. B. (2011). Heat avoidance is regulated by transient receptor potential (trp) channels and a neuropeptide signaling pathway in caenorhabditis elegans. Genetics, 188(1):91–103.

Hassold, T. and Hunt, P. (2001). To err (meiotically) is human: the genesis of human aneuploidy. Nature Reviews Genetics, 2(4):280–291.

Hill, W. G. and Robertson, A. (1966). The effect of linkage on limits to artificial selection. Genetics Research, 8(3):269–294.

Hillers, K. J. and Villeneuve, A. M. (2003). Chromosome-wide control of meiotic crossing over in c. elegans. Current Biology, 13(18):1641–1647.

Hodgkin, J. and Doniach, T. (1997). Natural variation and copulatory plug formation in caenorhabditis elegans. Genetics, 146(1):149–164.

Hodgkin, J., Horvitz, H. R., and Brenner, S. (1979). Nondisjunction mutants of the nematode caenorhab-ditis elegans. Genetics, 91(1):67–94.

i Torres, A. P., Höök, L., Näsvall, K., Shipilina, D., Wiklund, C., Vila, R., Pruisscher, P., and Backström, N. (2023). The fine-scale recombination rate variation and associations with genomic features in a butterfly. Genome Research, 33(5):810–823.

Johnston, H. R. and Cutler, D. J. (2012). Population demographic history can cause the appearance of recombination hotspots. The American Journal of Human Genetics, 90(5):774–783.

Kulathinal, R. J., Bennett, S. M., Fitzpatrick, C. L., and Noor, M. A. (2008). Fine-scale mapping of recombination rate in drosophila refines its correlation to diversity and divergence. Proceedings of the National Academy of Sciences, 105(29):10051–10056.

Lam, I. and Keeney, S. (2015). Mechanism and regulation of meiotic recombination initiation. Cold Spring Harbor perspectives in biology, 7(1):a016634.

Lenormand, T. and Otto, S. P. (2000). The evolution of recombination in a heterogeneous environment. Genetics, 156(1):423–438.

Lim, J. G., Stine, R. R., and Yanowitz, J. L. (2008). Domain-specific regulation of recombination in caenorhabditis elegans in response to temperature, age and sex. Genetics, 180(2):715–726.

McGrath, P. T., Rockman, M. V., Zimmer, M., Jang, H., Macosko, E. Z., Kruglyak, L., and Bargmann, C. I. (2009). Quantitative mapping of a digenic behavioral trait implicates globin variation in c. elegans sensory behaviors. Neuron, 61(5):692–699.

McVean, G. A., Myers, S. R., Hunt, S., Deloukas, P., Bentley, D. R., and Donnelly, P. (2004). The fine-scale structure of recombination rate variation in the human genome. Science, 304(5670):581–584.

Meneely, P. M., Farago, A. F., and Kauffman, T. M. (2002). Crossover distribution and high interference for both the x chromosome and an autosome during oogenesis and spermatogenesis in caenorhabditis elegans. Genetics, 162(3):1169–1177.

Noble, L. M., Yuen, J., Stevens, L., Moya, N., Persaud, R., Moscatelli, M., Jackson, J. L., Zhang, G., Chitrakar, R., Baugh, L. R., et al. (2021). Selfing is the safest sex for caenorhabditis tropicalis. Elife, 10:e62587.

Otto, S. P. (2009). The evolutionary enigma of sex. the american naturalist, 174(S1):S1–S14.

Otto, S. P. and Barton, N. H. (2001). Selection for recombination in small populations. Evolution, 55(10):1921–1931.

Paigen, K. and Petkov, P. M. (2018). Prdm9 and its role in genetic recombination. Trends in Genetics, 34(4):291–300.

Parée, T. and Teotónio, H. (2025). Experimental tests on the evolution of sex and recombination and their adaptive significance. Journal of Evolutionary Biology, 38(7):798–810.

Ptak, S. E., Hinds, D. A., Koehler, K., Nickel, B., Patil, N., Ballinger, D. G., Przeworski, M., Frazer, K. A., and Pääbo, S. (2005). Fine-scale recombination patterns differ between chimpanzees and humans. Nature genetics, 37(4):429–434.

Rockman, M. V. and Kruglyak, L. (2009). Recombinational landscape and population genomics of caenorhabditis elegans. PLoS genetics, 5(3):e1000419.

Rose, A. and Baillie, D. (1979). The effect of temperature and parental age on recombination and nondisjunction in caenorhabditis elegans. Genetics, 92(2):409–418.

Schedl, T. and Kimble, J. (1988). fog-2, a germ-line-specific sex determination gene required for hermaphrodite spermatogenesis in caenorhabditis elegans. Genetics, 119(1):43–61.

Shakes, D. C., Wu, J.-c., Sadler, P. L., LaPrade, K., Moore, L. L., Noritake, A., and Chu, D. S. (2009). Spermatogenesis-specific features of the meiotic program in caenorhabditis elegans. PLoS genetics, 5(8):e1000611.

Singhal, S., Leffler, E. M., Sannareddy, K., Turner, I., Venn, O., Hooper, D. M., Strand, A. I., Li, Q., Raney, B., Balakrishnan, C. N., et al. (2015). Stable recombination hotspots in birds. Science, 350(6263):928–932.

Sterken, M. G., Snoek, L. B., Kammenga, J. E., and Andersen, E. C. (2015). The laboratory domestication of caenorhabditis elegans. Trends in Genetics, 31(5):224–231.

Stevison, L. S., Woerner, A. E., Kidd, J. M., Kelley, J. L., Veeramah, K. R., McManus, K. F., Project, G. A. G., Bustamante, C. D., Hammer, M. F., and Wall, J. D. (2016). The time scale of recombination rate evolution in great apes. Molecular biology and evolution, 33(4):928–945.

Stiernagle, T. (2006). Maintenance of c. elegans. WormBook: The online review of C. elegans biology [Internet].

Stiernagle, T. et al. (1999). Maintenance of c. elegans.

Styer, K. L., Singh, V., Macosko, E., Steele, S. E., Bargmann, C. I., and Aballay, A. (2008). Innate immunity in caenorhabditis elegans is regulated by neurons expressing npr-1/gpcr. Science, 322(5900):460–464.

Teterina, A. A., Willis, J. H., Lukac, M., Jovelin, R., Cutter, A. D., and Phillips, P. C. (2023). Genomic diversity landscapes in outcrossing and selfing caenorhabditis nematodes. PLoS Genetics, 19(8):e1010879.

Thompson, O. A., Snoek, L. B., Nijveen, H., et al. (2015). Remarkably divergent regions punctuate the genome assembly of the caenorhabditis elegans hawaiian strain cb4856. Genetics.

Tsai, C. J., Mets, D. G., Albrecht, M. R., Nix, P., Chan, A., and Meyer, B. J. (2008). Meiotic crossover number and distribution are regulated by a dosage compensation protein that resembles a condensin subunit. Genes & development, 22(2):194–211.

Volkers, R. J., Snoek, L. B., Hubar, C. J. v. H., Coopman, R., Chen, W., Yang, W., Sterken, M. G., Schulenburg, H., Braeckman, B. P., and Kammenga, J. E. (2013). Gene-environment and protein-degradation signatures characterize genomic and phenotypic diversity in wild caenorhabditis elegans populations. BMC biology, 11(1):93.

Weber, K. P., De, S., Kozarewa, I., Turner, D. J., Babu, M. M., and de Bono, M. (2010). Whole genome sequencing highlights genetic changes associated with laboratory domestication of c. elegans. PloS one, 5(11):e13922.

Zetka, M.-C. and Rose, A. M. (1990). Sex-related differences in crossing over in caenorhabditis elegans. Genetics, 126(2):355–363.

Zhao, Y., Long, L., Xu, W., Campbell, R. F., Large, E. E., Greene, J. S., and McGrath, P. T. (2018). Changes to social feeding behaviors are not sufficient for fitness gains of the caenorhabditis elegans n2 reference strain. Elife, 7:e38675.

