## Supplemental figures and methods for "Inter-strain variation in intra-chromosomal rates of recombination in *Caenorhabditis elegans*"

Supplemental Files

Supplemental Figures

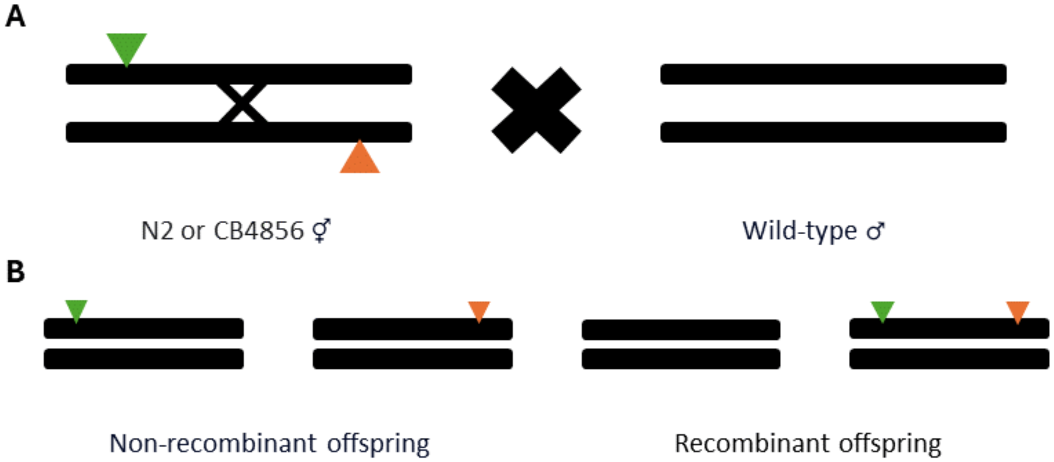


Figure 1: The crossing designed to estimate recombination rate of heterozygous hermaphrodite with two different markers, between hermaphrodite and wild-type male. **A)** The focal individual is doubly heterozygous, with markers on different copies of chromosome IV. The focal individual is crossed to a wild-type mate that do not carry any marker alleles. **B)** There are four possible phenotypes of offspring that can result from this cross: two non-recombination phenotypes (offspring with only a GFP marker or a tdTomato marker) and two recombinant phenotypes (offspring with both markers and no markers). By counting the frequencies of each we can estimate r, the recombination rate between the two fluorescent markers. The probability of recombinant offspring is 0.5*r and the probability of non-recombinant offspring is 0.5*(1-r).

*Supplemental Methods*

*Inserting the fog-2 Mutation into N2/CB4856 Fluorescent Strains*

First, we generated heterozygous F1s by crossing the fluorescent strain and the *fog-2* strain (same N2 and CB4856 genetic background). Here we used hermaphrodites from the *fog-2* strain and males from the fluorescent strain to ensure outcrossing. Then, we crossed F1s to generate F2s. We selected 200 F2 fluorescent hermaphrodites at L4 stage and put them on individual plates. We discarded all of the non-fluorescent worms. We retained male F2s on the original plate to use for future crosses. We kept the fluorescent F2 hermaphrodites on the plates for 24 hours and observed if they produced viable offspring. We took this step to ensure that the hermaphrodites were homozygous for the *fog-2* gene and were not capable of producing embryos. We selected the plates with hermaphrodites that did not produce viable offspring and discarded the rest. Then, we crossed the selected hermaphrodites with fluorescent F2 males and obtained an F3 generation. Once the F3 offspring reached the L4 stage, we again examined each plate for fluorescence and discarded all of the plates with non-fluorescent worms. Then, from each of the remaining plates, we transferred 10 hermaphrodites to individual plates. We referred to the groups of 10 plates as batches. We again retained all of the F3 males on the original plates. We kept the hermaphrodites of these batches on the plates for 24 hours again and observed if they produced viable offspring. We selected two plates from each remaining batch that did not have embryos on the plates and crossed the hermaphrodites with an F3 male from their original plate. Lastly, we checked for fluorescence and only retained the plates with 100% homozygous fluorescent worms. We then maintained these fluorescent *fog-2* strains and used them to estimate the recombination rates of hermaphrodites and males.

*N2 and CB4856 Strains - Genetic Background Calculations*

This percentage was calculated by adding the percentage of genetic background offspring received from the parents for 10 generations. For instance, we start with a cross between an individual that carries a fluorescent marker and an individual that carries our desired genetic background (N2 or CB4856). From this cross, we obtain an F1 individual that is 50% of our desired genetic background. When we backcross the F1 individual we can expect the F2 individuals to be 75% of the desired genetic background. Similarly, when the F2 generation is backcrossed we can expect F3 offspring to be 87.5% of the desired genetic background. Therefore, after 10 rounds of backcrossing we can expect 99.951% of the desired genetic background with the focal fluorescent markers.

*Fluorescent Marker Distance*

We utilized reference genomes of N2 and CB4856 chromosome IV to determine if the distance between the fluorescent markers is similar in both strains and if it is not impacting the differences in recombination rates between strains. We utilized the flanking sequences given in the wormbuilder.org for strains with mCherry, GFP and tdTomato markers (Frøkjær-Jensen *et al.* 2014). We then used NCBI's BLAST to locate the exact nucleotide positions for the flanking sequences on N2 and CB4856 chromosome IV reference (Altschul *et al.* 1990). For the gene-poor region, once we acquired the exact nucleotide positions for both flanking sequences, we calculated the approximate distance between the first nucleotide position of the GFP flanking sequence to the first nucleotide of the tdTomato flanking position. We performed this calculation for both N2 and CB4856 strains. For the gene-rich center, we used the same methodology to calculate the distance between mCherry and GFP markers using their flanking sequences in the N2 strain. However, we could not locate the flanking sequence for mCherry marker on the CB4856 chromosome IV reference. To get an approximate distance for the marker location in CB4856 we used the Y46C8AL.12 gene that is right upstream and the closest to the mCherry marker (Howe *et al.* 2017). We then used BLAST to obtain the position of the gene on both N2 and CB4856 chromosome references. Once we obtained this, we calculated the distance between the first nucleotide position of the flanking sequence and the first nucleotide position of the Y46C8AL.12 gene on the N2 chromosome IV reference to obtain an approximate distance between the mCherry marker and the Y46C8AL.12 gene (1007 bp). We then used this approximate distance (1007 bp) to locate where the mCherry marker flank would be on the CB4856 chromosome IV reference by subtracting the distance between the markers (1007 bp) from the first nucleotide position of the Y46C8AL.12 gene of the CB4856 chromosome IV. Once we obtained the position where the mCherry marker flank would be on chromosome IV of the CB4856 strain, we calculated the distance between mCherry and GFP marker flanks to get the approximate distance between the focal markers in the CB4856 strain.

### References

Altschul SF, Gish W, Miller W, Myers EW, Lipman DJ. 1990. Basic local alignment search tool. Journal of molecular biology. 215:403–410.

Frøkjær-Jensen C, Davis MW, Sarov M, Taylor J, Flibotte S, LaBella M, Pozniakovsky A, Moerman DG, Jorgensen EM. 2014. Random and targeted transgene insertion in caenorhabditis elegans using a modified mos1 transposon. Nature methods. 11:529–534.

Howe KL, Bolt BJ, Shafie M, Kersey P, Berriman M. 2017. Wormbase parasite- a comprehensive resource for helminth genomics. Molecular and biochemical parasitology. 215:2–10.
